# Macropinocytosis of amyloid precursor protein requires the adaptor protein Fe65 and the recruitment and activity of Arf6 and the RhoGTPases Rac1, Cdc42 and RhoA

**DOI:** 10.1101/2025.03.02.641070

**Authors:** Jordan M. Krupa, Abdul M. Naqvi, Manoj Reddy Medapati, Adrianna R. Tsang, Claudia Seah, Stephen H. Pasternak

## Abstract

Alzheimer’s disease (AD) is a progressive neurodegenerative disorder characterized by the buildup of the highly toxic peptide amyloid-beta (Aβ). Previously, we demonstrated that Aβ is generated from the cleavage of amyloid precursor protein (APP) after internalization to lysosomes via macropinocytosis. However, the regulation of APP micropinocytosis has remained uncharacterized. Evidence suggests that APP may function as a cell surface receptor which could contribute to this regulation. Arf6 and the RhoGTPases Rac1, Cdc42 and RhoA are known to regulate macropinocytosis in response to signaling of other receptors. An adaptor protein called Fe65, which can associate with both amyloid precursor protein and Arf6, could function as the link between APP and these known regulatory elements. Thus, we hypothesized that the binding and/or crosslinking of APP recruits Fe65, which then recruits and activates Arf6, which in turn activates Rac1, Cdc42 and RhoA, resulting in APP macropinocytosis. Rapid and transient recruitment of Fe65 and Arft6 was observed to APP 30 seconds following binding/crosslinking. Rac1, Cdc42 and RhoA all examined demonstrated more sustained recruitment to crosslinked APP. Prevention of Fe65 binding by APP mutation and Arf6 inhibition by NAV-2729 prevented the recruitment of all proteins. Together, these observations are the first to demonstrate that a network of regulatory proteins is recruited to bound/crosslinked APP which regulates its macropinocytosis. Targeting these regulatory proteins could be explored to modulate the membrane to lysosomal trafficking of APP and reducing the production of Aβ in AD.

## Introduction

Alzheimer’s disease (AD) is a progressive neurodegenerative disorder that affects roughly 43.8 million people worldwide, a number that is expected to significantly increase to 152 million by 2050 [1]. AD is characterized on a cellular level by the formation of amyloid plaques containing the protein amyloid-β (Aβ), and neurofibrillary tangles composed of the protein tau [2,3]. Aβ peptides and soluble aggregates have been found to be highly toxic to neurons and to dendritic spines and synapses, where synaptic loss is strongly correlated with impaired cognition in AD [4]. Moreover, Aβ can seed tau pathology, which is more closely related to neuronal cell death [5].

The Aβ peptide is produced by the enzymatic cleavage of the type-I transmembrane protein amyloid precursor protein (APP) by β-secretase (BACE) and γ-secretase [2,3]. Amyloidogenic processing of APP begins with BACE cleaving off a N-terminal portion of APP (sβAPP), followed by γ-secretase cleavage within the lipid bilayer to release the toxic Aβ peptide and APP intracellular domain (AICD) [3]. The production of the Aβ peptide is followed by its either clearance or aggregation into soluble and Aβ oligomers, which can further aggregate to form insoluble aggregates [3]. The exact sub-cellular site of the amyloidogenic cleavage of APP has been hard to identify, but evidence suggests the lysosome plays a critical role. This is supported by early observations that intracellular Aβ colocalizes with autophagosomes and lysosomes [6,7]. More importantly, lysosomal membranes are enriched in APP, the BACE cleavage product β-C-terminal fragment (βCTF), and the components of γ-secretase [8–10]. Additionally, lysosomes have been demonstrated to have an acidic pH optimal to γ-secretase functioning [11] and inhibition of γ-secretase has been shown to cause amyloidogenic fragments of APP to accumulate in lysosomes [12].

Previously, we identified a novel pathway that trafficked APP rapidly to lysosomes [13], which we revealed to be macropinocytosis [14]. Macropinocytosis is an endocytic pathway that internalizes large amounts of plasma membrane and extracellular fluid [15]. It is mediated by actin polymerization at the plasma membrane, which generates membrane ruffles, which can then fuse onto themselves to form macropinocytic cups. Cups can then collapse on the plasma membrane forming a macropinosomes, which rapidly fuse with lysosomes near the plasma membrane [16]. Throughout these distinct stages of macropinocytosis, membrane phosphatidylinositol (PI) undergoes specific changes to its phosphorylation state [17,18]. Membrane ruffling in macropinocytosis has been demonstrated to be enriched in PI(4,5)P_2_ and can be visualized by fluorescently tagging the PH domain from PLC8 (PLC8PH) [19,20].

Although little is known about macropinocytosis in neuronal cells, in other cell types it has been demonstrated to be stimulated by receptor binding, for example by signaling produced by growth factor receptors [21]. APP could behave similarly as it contains ligand binding domains [22], and the binding and/or crosslinking of APP using an antibody has been shown to result in increased internalization and Aβ production [14,23]. Previously, we demonstrated that the internalization of cell surface APP to lysosomes occurs constitutively and is significantly increased by antibody binding [14]. Several small GTPases have been implicated in the regulation of macropinocytosis in non-neuronal cells [21]. The regulatory small GTPase Arf6 is known to be involved in the initiation of macropinocytosis [21,24]. In regards to APP, knockdown or dominant negative mutations in Arf6 have been demonstrated to reduce the amount of APP located in lysosomes and Aβ production [14]. In brain samples from human AD patients, increased Arf6 expression has been demonstrated in hippocampal regions, suggesting that this GTPase may be important in pathological Aβ production [14].

The Rho family of GTPases (RhoGTPases) including Rac1, Cdc42, and RhoA are involved in the actin polymerization and myosin light chain remodeling required for initiation of macropinocytosis [16,21]. Arf6 has been demonstrated to bind to and activate both Rac1 and Cdc42 through a binding partner [25]. Rac1 is involved in the formation of the membrane ruffles and lamellipodia [24], while Cdc42 activation results in the generation of filipodia and the macropinocytic cup [21,26]. Further, Rac1 had been previously demonstrated to activate RhoA, which is important for the formation of stress fibers [27]. All together, these changes are believed to link the plasma membrane to the actin cytoskeleton and initiate macropinocytosis [21]. There has been some evidence to implicate these proteins in AD. Knock down or dominant negative mutations in Rac1 have been shown to decrease the amount of APP located in lysosomes [14]. Additionally, inhibiting either Rac1 or RhoA has been demonstrated to reduce the production of Aβ [28,29].

Considering there are no current direct links between Arf6 and APP in the literature. An adaptor protein called Fe65 is known to be expressed in neurons and bind to the intracellular C-terminal YENPTY ‘endocytosis’ signal [30] as well as to Arf6, activating both Arf6 and Rac1 [31,32]. The amount of Fe65 in the CA4 region of the hippocampus is associated with the severity of AD [33]. Here, we hypothesize that binding and/or crosslinking cell surface APP results in the recruitment and binding of Fe65, which in turn recruits and activates Arf6, which then recruits and activates the RhoGTPases Rac1, RhoA and Cdc42, resulting in the macropinocytosis of APP to lysosomes.

Following APP labeling with a fluorescently labeled antibody, we observed rapid and transient recruitment of Fe65, Arf6 and each of the RhoGTPases examined to crosslinked APP and membrane ruffles. The recruitment of Rac1, Cdc42 and RhoA was longer lasting than either Fe65 or Arf6, suggesting they may play sustained roles downstream of these proteins. Further supporting their downstream role, mutation of APP to prevent Fe65 binding or pharmacological inhibition of Arf6 significantly reduced the recruitment of Rac1, Cdc42 and RhoA. To support the notion that increased recruitment coincides with increased activity of these GTPases, antibody binding of APP was observed to rapidly increase the amount of GTP-bound Rac1, Cdc42 and RhoA. Together, these results provide evidence for the existence of a regulatory cascade that is directly recruited to the plasma membrane at sites of bound/crosslinked APP, which regulates its trafficking to lysosomes through macropinocytosis. Given these small GTPases are readily targeted by small molecule inhibitors, the signaling cascade elucidated in this study could be targeted to reduce the production of Aβ peptides through modulation of APP trafficking to lysosomes.

## Results

### N-terminal APP antibody binding/crosslinking of cell surface APP drives internalization to lysosomes through macropinocytosis

Our previous studies examining APP macropinocytosis utilized anti-HA or 6E10 anti-APP antibodies to bind/crosslink HA-tagged APP or untagged APP695, respectively [13,14,34]. Here, we utilized an anti-APP antibody targeting an epitope containing amino acids 104-118 of APP, corresponding to the growth-factor like domain (GFLD) of the N-terminal E1 domain [35]. This antibody was chosen given macropinocytosis is frequently observed in response to growth factor ligand-receptor binding [36], and thus may represent a more physiologically relevant target. We first compared this antibody with 6E10, an anti-APP/Aβ antibody that we used in our previous work [13,14,34]. Neuro2a (N2a) cells were chosen as the model for these experiments as they are readily transduced by transfection, have been previously used for studies investigating protein GTP binding, and widely used in neurodegenerative research. N2a cells were transduced with APP695 and LAMP1-mCherry (LAMP1-mCh) were incubated with fluorescently labeled N-terminal APP antibody, 6E10 antibody, or NCAM antibody (negative control) on ice for 20 minutes, and then either fixed immediately (Time 0) or after incubation at 37 °C for 15 minutes. Cells were imaged using Laser Scanning Confocal Microscopy on a Leica SP8. Imaris 10.1 Software (Bitplane) was used to quantitate colocalization between markers and to generate a co-localization channel (white). At Time 0, each fluorescently tagged antibody was observed entirely around the plasma membrane (Figure 1A,C). After 15 minutes, both the N-terminal APP-targeting antibody (Figure 1A) and 6E10 antibody (Figure 1B) were observed colocalized with LAMP1 compartments, while NCAM remained at the cell surface (Figure 1C). When we quantitate the colocalization of labelled antibodies and LAMP1, antibodies targeting the N-terminal portion of APP or residues within the Aβ region (6E10) demonstrated significantly higher colocalization between tagged antibody and LAMP1 compared to antibodies directed towards NCAM after 15 minutes (Figure 1D; p<0.001). This supports that the N-terminal APP antibody targeting the GFLD also rapidly directs APP to lysosomes. To determine if this rapid uptake was due to macropinocytosis, uptake of bound/crosslinked APP to lysosomes was examined in cells treated with EIPA (macropinocytosis inhibitor) or Pitstop2 (clathrin-coat/CME inhibitor). Antibody labeled APP remains at the plasma membrane with minimal colocalization with LAMP1 after 15 minutes when treated with EIPA, suggesting its internalization by macropinocytosis has been prevented (Figure 1E). Internalization of antibody labeled APP to lysosomes can be clearly seen by the colocalization observed between APP and LAMP1 with Pitstop2 performing similar to DMSO treatment (Figure 1E). Comparing treatments, colocalization of APP with LAMP1 was significantly lower in EIPA treated cells compared to vehicle control or treatment with Pitstop2 (p<0.001; Figure 1F).

**Figure 1.**
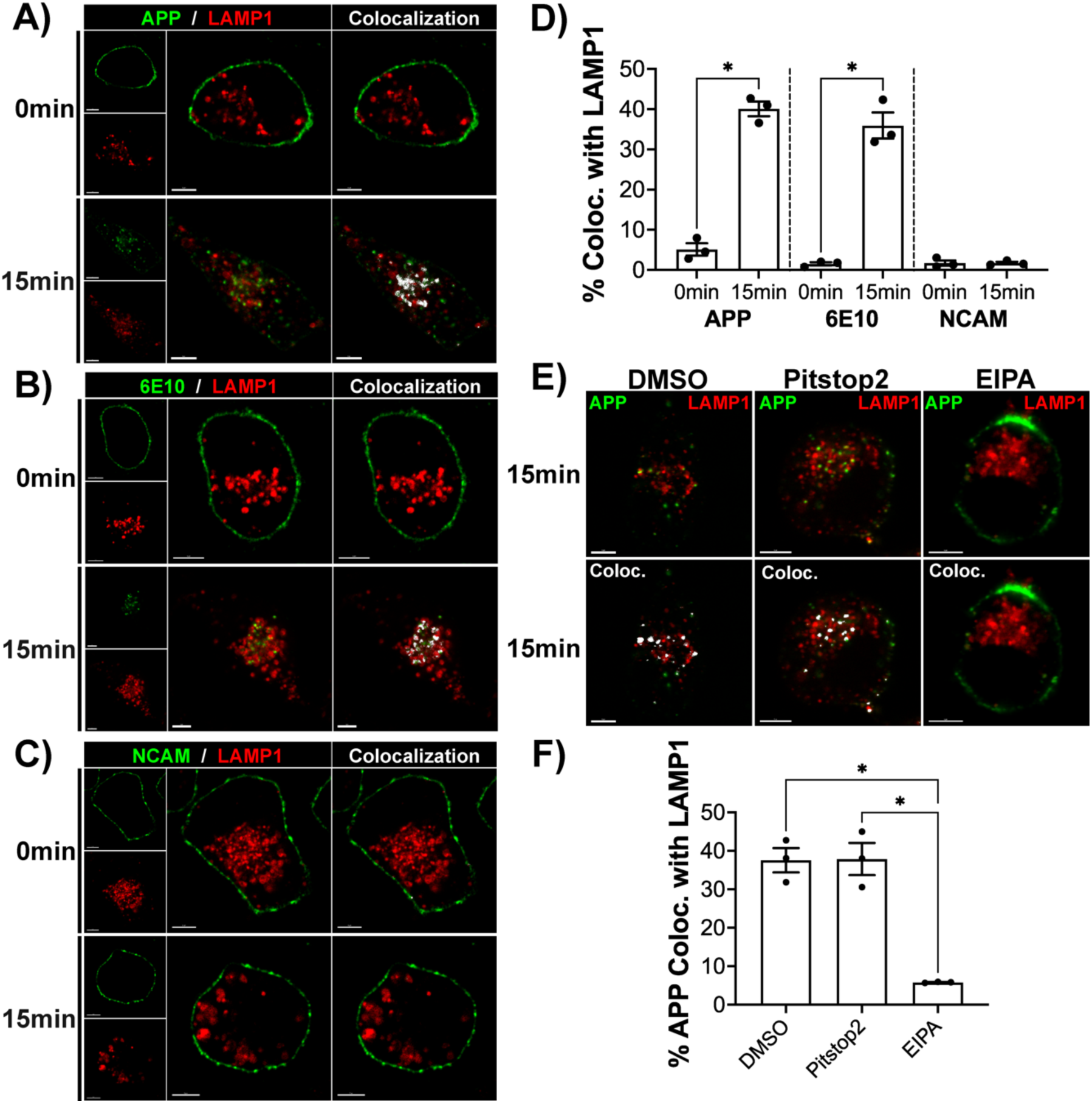
N-terminal anti-APP antibody mediated binding/crosslinking of APP results in rapid internalization to lysosomes by macropinocytosis. N2a cells transfected with APP695 and LAMP1-mCh (red) and incubated with fluorescently-tagged **A)** N-terminal anti-APP (APP), **B)** Aβ-region targeting 6E10 antibodies, or **C)** anti-NCAM antibodies (green) on ice for 20 minutes, then immediately fixed or allowed to incubate at 37°C for 15 minutes. Colocalization was assessed between crosslinked APP and LAMP1 (white pixels). **D)** Quantification of the mean % of APP colocalized with LAMP1 from three replicate experiments (n=3; 10 images per replicate), with significance calculated by a one-way ANOVA with Tukey’s test. **E)** N2a cells transfected with APP695 and LAMP1-mCh (red) that were treated with DMSO, 20μM Pitstop2 or 10μM EIPA. APP was then bound/crosslinked by antibody and imaged after 15 minutes. Colocalization was assessed between crosslinked APP and LAMP1 (white pixels). **F)** Quantification of the mean % of APP colocalized with LAMP1 (n=3; 10 images per replicate), with significance calculated by a one-way ANOVA with Tukey’s test. *Data is presented as mean ± SEM. * p<0.05; Scale bar = 5μm*.

### Fe65 and Arf6 are rapidly and transiently recruited to bound/crosslinked APP and membrane ruffles

This antibody-mediated APP binding/crosslinking experiment was then utilized to assess the recruitment of Fe65 to crosslinked APP (Figure 2A). N2a cells transduced with APP695, Fe65-EGFP and the ruffling membrane marker PLC8PH-mCherry were incubated with fluorescently labeled APP or NCAM antibodies on ice for 20 minutes, and then either fixed immediately (Time 0) or after incubation at 37°C for the times indicated. Imaris 10.1 Software (Bitplane) was used to quantitate colocalization and generate a co-localization channel (white), which was overlayed to highlight colocalization. In response to the antibody-mediated binding/crosslinking of APP, Fe65 rapidly colocalized with APP within 30 seconds following removal from ice, demonstrated by the colocalization observed between the two at that timepoint (Figure 2A). Whereas no discernable recruitment was observed in response to NCAM binding/crosslinking (Figure S1A). Additionally, this rapid recruitment occurred in membrane ruffles marked by PLC8PH, demonstrated by the colocalization between Fe65 and PLC8PH (Figure 2A). Significantly larger differences in percent colocalized were demonstrated at 30 seconds between Fe65 and bound/crosslinked APP (8.68 ± 0.484; p<0.001; Figure 2B) or PLC8PH (9.16 ± 0.292; p<0.001; Figure 2C) compared to baseline. When NCAM was bound/crosslinked, there were no significant increases or decreases at any timepoint compared to baseline (Figure 2B,C). Comparing APP to NCAM binding/crosslinking at each time point, significantly higher difference to baseline was observed at 30 seconds for both colocalization measurements (p<0.001; Figure 2B,C).

**Figure 2.**
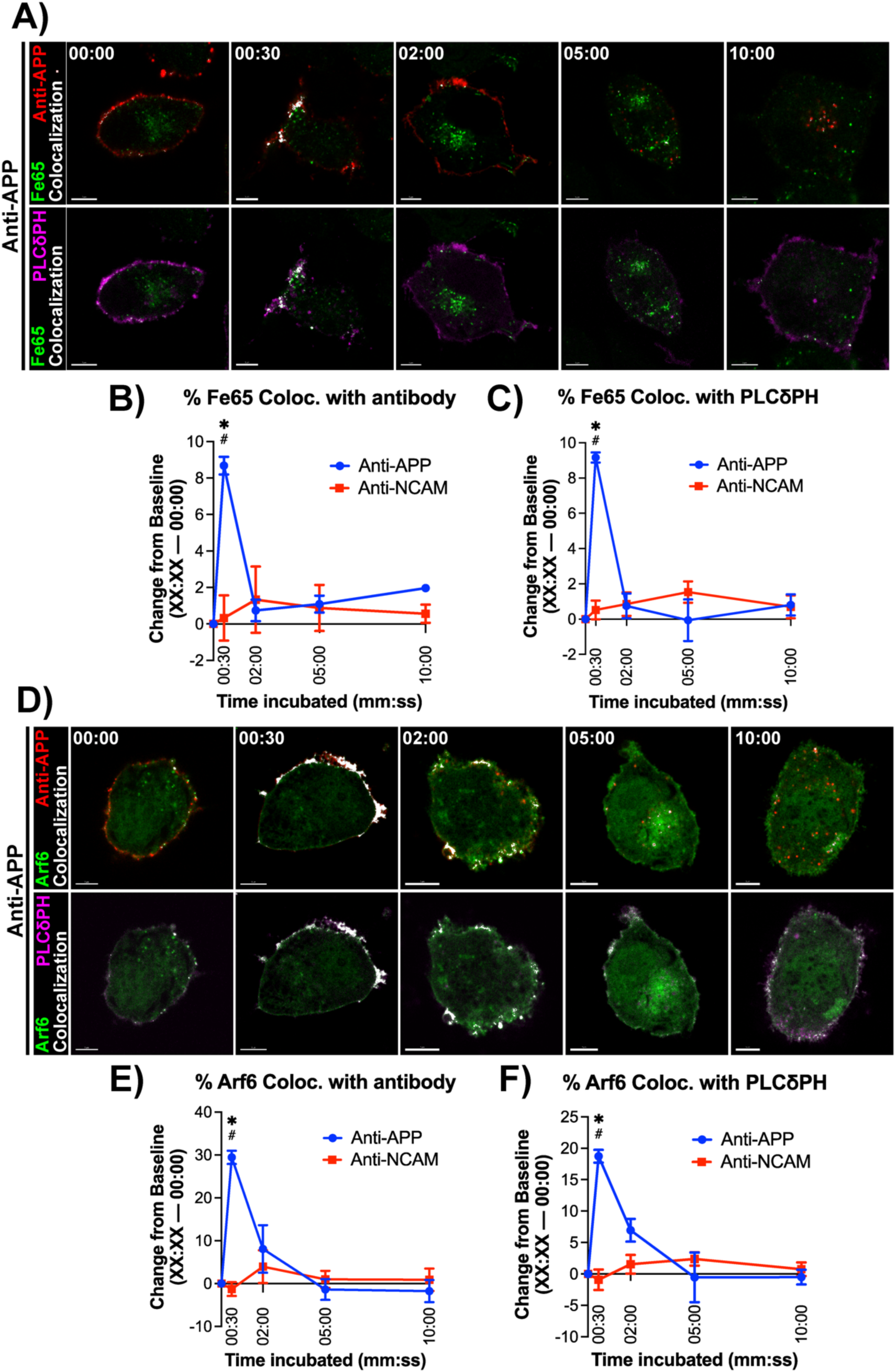
Fe65 and Arf6 are rapidly and transiently recruited to antibody bound/crosslinked APP and membrane ruffles after 30 seconds. **A)** N2a cells transfected with Fe65-EGFP (green), PLC8PH-mRFP (magenta) and APP695. Cells were incubated with fluorescent anti-APP or anti-NCAM antibodies (red) on ice, then immediately fixed on ice as a baseline (00:00) or incubated for 30 seconds (00:30), 2 minutes (02:00), 5 minutes (5:00), and 10 minutes (10:00). Colocalization between signals is indicated by white pixels. **B/C)** Quantification of colocalization data (n=3; 15 images per replicate) is represented as the difference between the colocalization at each timepoint to the colocalization at baseline (mean % colocalized at XX:XX – mean % colocalized at 00:00). * denotes significant difference to baseline within condition; # denotes significant difference between groups at the indicated timepoint. **D)** N2a cells transfected with Arf6-EGFP (green), PLC8PH-mRFP (magenta) and APP695. Experiment above was repeated with Arf6-EGFP expressing cells. Colocalization between signals is indicated by white pixels. **E/F)** Quantification of colocalization data (n=3; 15 images per replicate). Data representation and analysis from G/H) was repeated for N2a cells expressing Arf6-GFP. * denotes significant difference to baseline within condition; # denotes significant difference between groups at the indicated timepoint. *Significance was measured by a two-way ANOVA with a Tukey’s test using a single pooled variance. Data is presented as mean ± SEM. */# p<0.05; Scale bar = 5μm*.

Recruitment of the small GTPase Arf6 to bound/crosslinked APP and membrane ruffles was examined next. As before, cells expressing APP695, Arf6-EGFP and the ruffling membrane marker PLC8PH-mCherry were incubated with fluorescently labeled APP or NCAM antibodies on ice for 20 minutes, and then either fixed immediately (Time 0) or after incubation at 37 °C for the times indicated. Recruitment of Arf6 demonstrated a pattern similar to Fe65, with a peak in colocalization between Arf6 and bound/crosslinked APP or membrane ruffles observed within 30 seconds (Figure 2D). No changes in the amount of colocalization between Arf6 and labeled antibody or membrane ruffles were observed in response to NCAM binding/crosslinking (Figure S1B). Significantly larger difference to baseline was observed at 30 seconds with APP binding/crosslinking when analyzing colocalization between Arf6 and anti-APP (29.4 ± 1.54; p<0.001; Figure 2E) and PLC8PH (18.7 ± 1.02; p<0.001; Figure 2F). No significant differences in mean difference to baseline were observed within anti-NCAM treated timepoints (Figure 2E,F). At 30 seconds, anti-APP treatment demonstrated a significantly larger difference to baseline than anti-NCAM treatment for Arf6 colocalization with tagged antibody (p<0.001; Figure 2E) or PLC8PH (p<0.001; Figure 2F).

### Rac1, Cdc42 and RhoA demonstrate rapid and sustained recruitment to bound/crosslinked APP and membrane ruffles

We next assessed the recruitment of the RhoGTPases Rac1, Cdc42 and RhoA as described above with Fe65 and Arf6. Cells expressing APP695, Rac1-EGFP and the ruffling membrane marker PLC8PH-mCherry were incubated with fluorescently labeled APP or NCAM antibodies on ice for 20 minutes, and then either fixed immediately (Time 0) or after incubation at 37 °C for the times indicated. Imaris 10.1 Software (Bitplane) was used to quantitate colocalization and generate a co-localization channel (white), which was overlayed to highlight colocalization. Colocalization of Rac1 was observed with bound/crosslinked APP and PLC8PH at 30 seconds (Figure 3A), as was observed with Fe65 and Arf6. However, colocalization continued to be observed after 2 minutes of incubation (Figure 3A). Binding/crosslinking of NCAM produced no visible changes in colocalization with either tagged antibody or membrane ruffles (Figure S1C). Significantly higher differences in the percentage of Rac1 colocalized with anti-APP antibody were observed at 30 seconds (29.44 ± 3.720; p<0.001) and 2 minutes (24.87 ± 3.720; p<0.001) compared to baseline (Figure 3B). The same was observed with the colocalization between Rac1 and PLC8PH at 30 seconds (26.65 ± 7.072; p=0.009) and 2 minutes (19.48 ± 7.072; p=0.05; Figure 3C).

**Figure 3.**
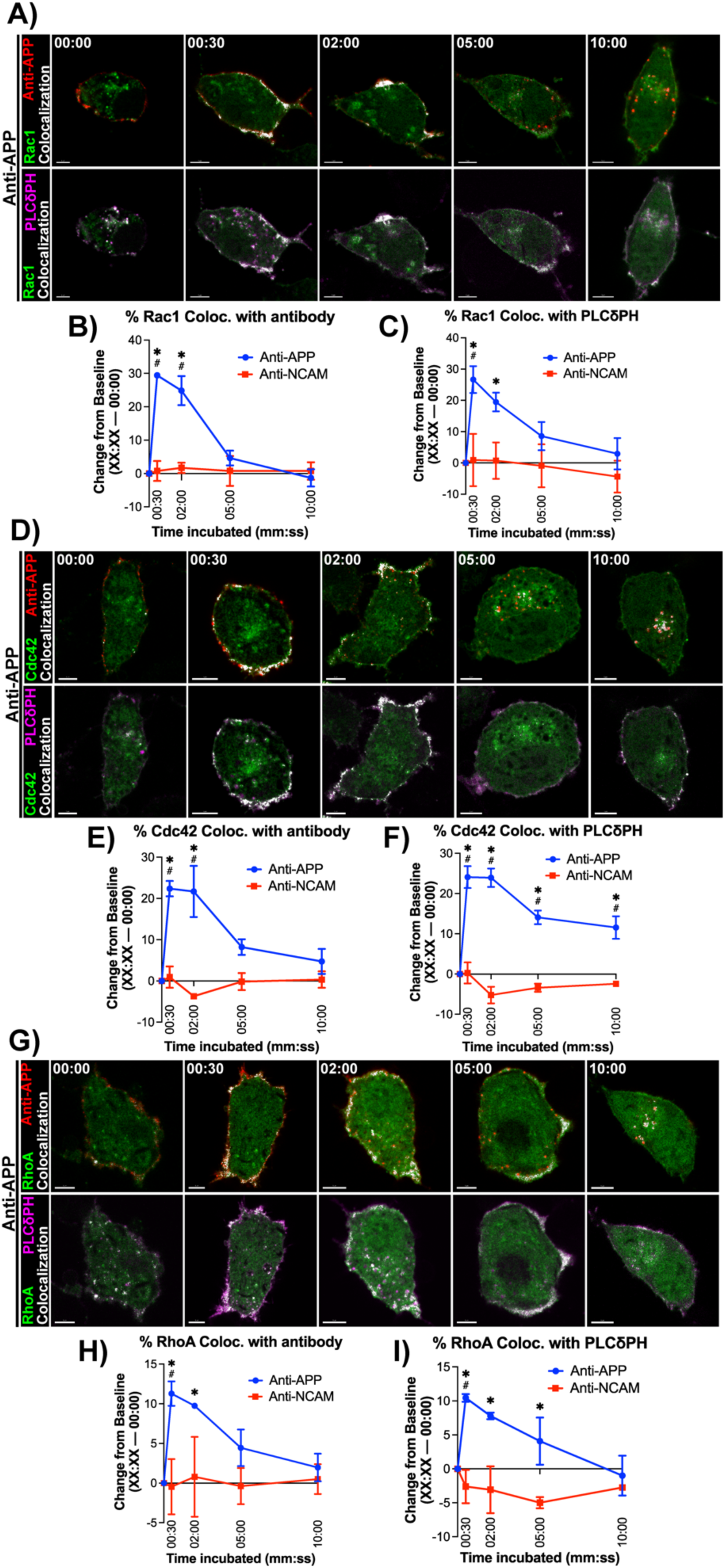
Rac1, Cdc42 and RhoA demonstrate rapid and sustained recruitment to bound/crosslinked APP and membrane ruffles. **A)** N2a cells transfected with Rac1-EGFP (green), PLC8PH-mRFP (magenta) and APP695. Cells were incubated with tagged anti-APP or anti-NCAM antibodies (red) on ice and then were either immediately fixed on ice as a baseline (00:00), or incubated for 30 seconds (00:30), 2 minutes (02:00), 5 minutes (5:00), and 10 minutes (10:00) at 37°C 5% CO2 prior to fixation. Colocalization between signals is indicated by white pixels. **B/C)** Quantification of colocalization data (n=3; 15 images per replicate) is represented as the difference between the colocalization at each timepoint to the colocalization at baseline (mean % colocalized at XX:XX – mean % colocalized at 00:00). * denotes significant difference to baseline within condition; # denotes significant difference between groups at the indicated timepoint. **D)** N2a cells transfected with Cdc42-EGFP (green), PLC8PH-mRFP (magenta) and APP695. Experiment from A) was repeated with Cdc42-EGFP expressing cells. Colocalization between signals is indicated by white pixels. **E/F)** Quantification of colocalization data (n=3; 15 images per replicate). Data representation and analysis from B/C) was repeated for N2a cells expressing Cdc42-EGFP. * denotes significant difference to baseline within condition; # denotes significant difference between groups at the indicated timepoint. **G)** N2a cells transfected with RhoA-EGFP (green), PLC8PH-mRFP (magenta) and APP695. Experiment from A) was repeated with RhoA-EGFP expressing cells. Colocalization between signals is indicated by white pixels. **H/I)** Quantification of colocalization data (n=3; 15 images per replicate). Data representation and analysis from B/C) was repeated for N2a cells expressing RhoA-EGFP. * denotes significant difference to baseline within condition; # denotes significant difference between groups at the indicated timepoint. *Significance was measured by a two-way ANOVA with a Tukey’s test using a single pooled variance. Data is presented as mean ± SEM. */# p<0.05; Scale bar = 5μm*.

Next cells expressing APP695, Cdc42-EGFP and the ruffling membrane marker PLC8PH-mCherry were incubated with fluorescently labeled APP, or NCAM antibodies on ice for 20 minutes, and then either fixed immediately (Time 0) or after incubation at 37°C for the times indicated. Imaris 10.1 Software (Bitplane) was used to quantitate colocalization and generate a co-localization channel (white), which was overlayed to highlight colocalization. Similar observations were made when examining the recruitment of Cdc42, with colocalization visually peaking at 30 seconds and 2 minutes following APP binding/crosslinking (Figure 3D), and no effect was on colocalization at any timepoint being observed in response to NCAM binding/crosslinking (Figure S1D). Significantly higher difference to baseline was observed in the colocalization between Cdc42 and bound/crosslinked APP at 30 seconds (22.4 ± 1.86; p<0.001) and 2 minutes (21.7 ± 1.89; p<0.001; Figure 3E). However, difference to baseline remained significantly increased between Cdc42 and PLC8PH at all timepoints (Figure 3F): 30 seconds (24.1 ± 2.72), 2 minutes (23.9 ± 2.29; p<0.001), 5 minutes (14.1 ± 1.70; p<0.001) and 10 minutes (11.6 ± 2.80; p<0.001). This may suggest that Cdc42 remained enriched at the ruffling membrane as the majority of APP was internalized into the cell and no longer present within membrane ruffles. No differences were observed between any timepoint when NCAM was bound/crosslinked, and difference to baseline was significantly higher for anti-APP treatment compared to anti-NCAM at 30 seconds, and 2 minutes for both colocalization measurements assessed (p<0.001; Figure 3E,F).

Finally, cells expressing APP695, RhoA-EGFP and the ruffling membrane marker PLC8PH-mCherry were incubated with fluorescently labeled APP or NCAM antibodies on ice for 20 minutes and then either fixed immediately (Time 0) or after incubation at 37°C for the times indicated. Similar observations to Rac1 and Cdc42 were also made with RhoA. Increased colocalization between both RhoA and bound/crosslinked APP or PLC8PH was seen in cells at 30 seconds and 2 minutes, with colocalization decreasing at 5 and 10 minutes (Figure 3G). No changes in colocalization over time were observed in response to the binding/crosslinking of NCAM (Figure S1E). Quantification of the difference in colocalization between each time point and baseline revealed a significant increase in RhoA – anti-APP colocalization from baseline at 30 seconds (11.3 ± 1.54; p=0.002) and 2 minutes (9.74 ± 0.342; p=0.02; Figure 3H). For colocalization of RhoA with PLC8PH, significantly higher difference to baseline was observed in response to APP binding/crosslinking at 30 seconds (10.4 ± 0.576; p<0.001), 2 minutes (7.80 ± 0.494; p<0.001) and 5 minutes (4.08 ± 3.46; p=0.004; Figure 3I). No significant increases or decreases were observed between either RhoA and antibody, or RhoA and PLC8PH in response to NCAM binding/crosslinking (Figure 3H,I). A significant difference between conditions was only observed at the 30 second timepoint for RhoA colocalization with antibody (p=0.02; Figure 3H) and PLC8PH (p=0.01; Figure 3I). Collectively, the RhoGTPases Rac1, Cdc42 and RhoA all demonstrated this sustained recruitment through 2 minutes, not observed with Fe65 or Arf6. This may indicate their activity is required in later aspects of macropinocytosis regulation and thus could serve downstream of Fe65 and Arf6 as part of a larger signalling cascade.

We also examined the activation of Rac1, Cdc42, and RhoA following antibody binding. To do so, cell expressing APP695 were incubated with APP or NCAM antibodies on ice for 20 minutes, and then incubated cells for 2 minutes at 37°C to allow the stimulation of APP macropinocytosis. Lysates were immediately collected from cells and were used to assess the levels of GTP-bound Rac1, Cdc42 and RhoA by G-LISA (Cytoskeleton). This experiment was also run in untreated cells, cells with incubated with anti-NCAM antibodies (negative control), and cells treated with EGF (positive control). Significantly higher absolute optical density was measured in response to both APP binding/crosslinking and positive control compared to either NCAM binding/crosslinking or untreated cells in Rac1 G-LISAs (p<0.001; Figure 4A), Cdc42 G-LISAs (p<0.001; Figure 4B), and RhoA GLISAs (p<0.001; Figure 4C). Increased absolute optical density measurements in response to antibody-mediated APP binding/crosslinking indicates higher levels of GTP-bound or active Rac1, Cdc42 and RhoA.

**Figure 4.**
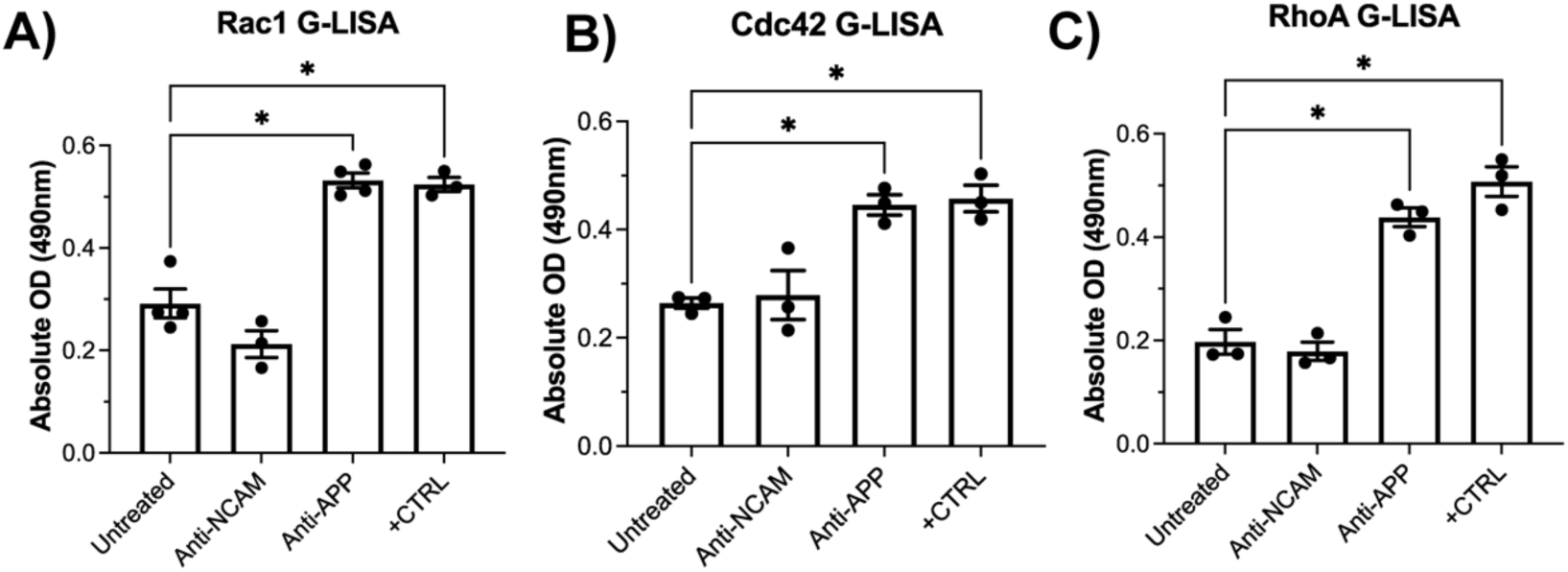
Antibody-mediated binding/crosslinking of APP increases GTP-bound Rac1, Cdc42 and RhoA levels. **A)** G-LISA assay for Rac1-GTP from N2a cells incubated with no treatment, anti-NCAM antibodies, anti-APP antibodies or EGF (positive control) while on ice, then incubated for 5 minutes to allow for the initiation of macropinocytosis. Lysates were immediately collected and used for Rac1 G-LISA activation assays. To compare the relative amount of active GTP-bound Rac1 in each condition, the absolute optical density (OD) was measured from each condition’s lysate loaded into G-LISA assay wells. Three different lysate samples were collected for each condition from different biological samples (n=3), and the G-LISA assay was performed in technical triplicates. Significant differences from untreated samples were calculated by a one-way ANOVA with Tukey’s test. This same experiment and analysis were performed for **B)** Cdc42-GTP levels, and **C)** RhoA-GTP levels. *Data is presented as mean* ± *SEM. *p<0.05*.

### Mutation of APP intracellular domain YENPTY sequence reduces recruitment of Fe65, Arf6, and the RhoGTPases Rac1, Cdc42 and RhoA

The C-terminal YENPTY sequence on APP has been shown to mediate the binding of Fe65 [37]. To examine the role of Fe65 in the macropinocytosis of APP to lysosomes and the recruitment of GTPases, N2a cells were transduced with APP696 containing either APP695 with the wildtype YENPTY sequence (WT-APP695) or had this sequence mutated to AENATA (APP695-AENATA) along with LAMP1-mCh and were incubated with fluorescently labeled APP antibodies on ice for 20 minutes, then allowed to internalize for 15 minutes at 37°C, fixed and imaged. This confirmed that APP-AENATA is not transported rapidly to lysosomes (Figure 5A,B). We then examined the effect of APP-AENATA on recruitment of each protein of interest after 30 seconds. Cells were transduced with either WT-APP or APP-AENATA and the ruffling membrane marker PLC8PH-mRFP along with each protein; Fe65-EGFP, Arf6-EGFP, Rac1-EGFP, or RhoA-EGFP. Transfected cells were incubated on ice with fluorescently labeled anti-APP antibodies for 20 minutes and then incubated for 30 seconds at 37°C, fixed and imaged. First, examining Fe65 recruitment, the binding/crosslinking of APP-AENATA resulted in significantly lower percent of Fe65 colocalized with both crosslinked APP (3.52% ± 0.674; p<0.001) and PLC8PH (6.84% ± 0.640; p<0.001) compared to WT-APP (Figure 5C,D). Arf6 recruitment was also reduced with mutation of the YENPTY sequence, with the cells transfected with APP-AENATA showing a significantly lower percent of Arf6 colocalized with bound/crosslinked APP (-13.6% ± 1.31; p<0.001) and PLC8PH (-19.8% ± 1.35; p=0.002) compared to WT (Figure 5E,F). This was also observed when examining the RhoGTPases Rac1, Cdc42 and RhoA. APP-AENATA expressing cells showed lower Rac1-APP colocalization (-14.9 ± 2.60; p=0.005) and Rac1-PLC8PH colocalization (-23.99 ± 3.137; p=0.002) than WT (Figure 5G,H). Percent colocalization of Cdc42 with APP (-22.31 ± 3.103; p=0.002; Figure 5I,J) or PLC8PH (-28.45 ± 3.385; p=0.005; Figure 4I,J), as well as RhoA with APP (-10.34 ± 2.187; p=0.009; Figure 4K,L) or PLC8PH (-14.61 ± 2.883; p=0.007; Figure 5K,L) was significantly lower as a result of crosslinking APP-AENATA.

**Figure 5.**
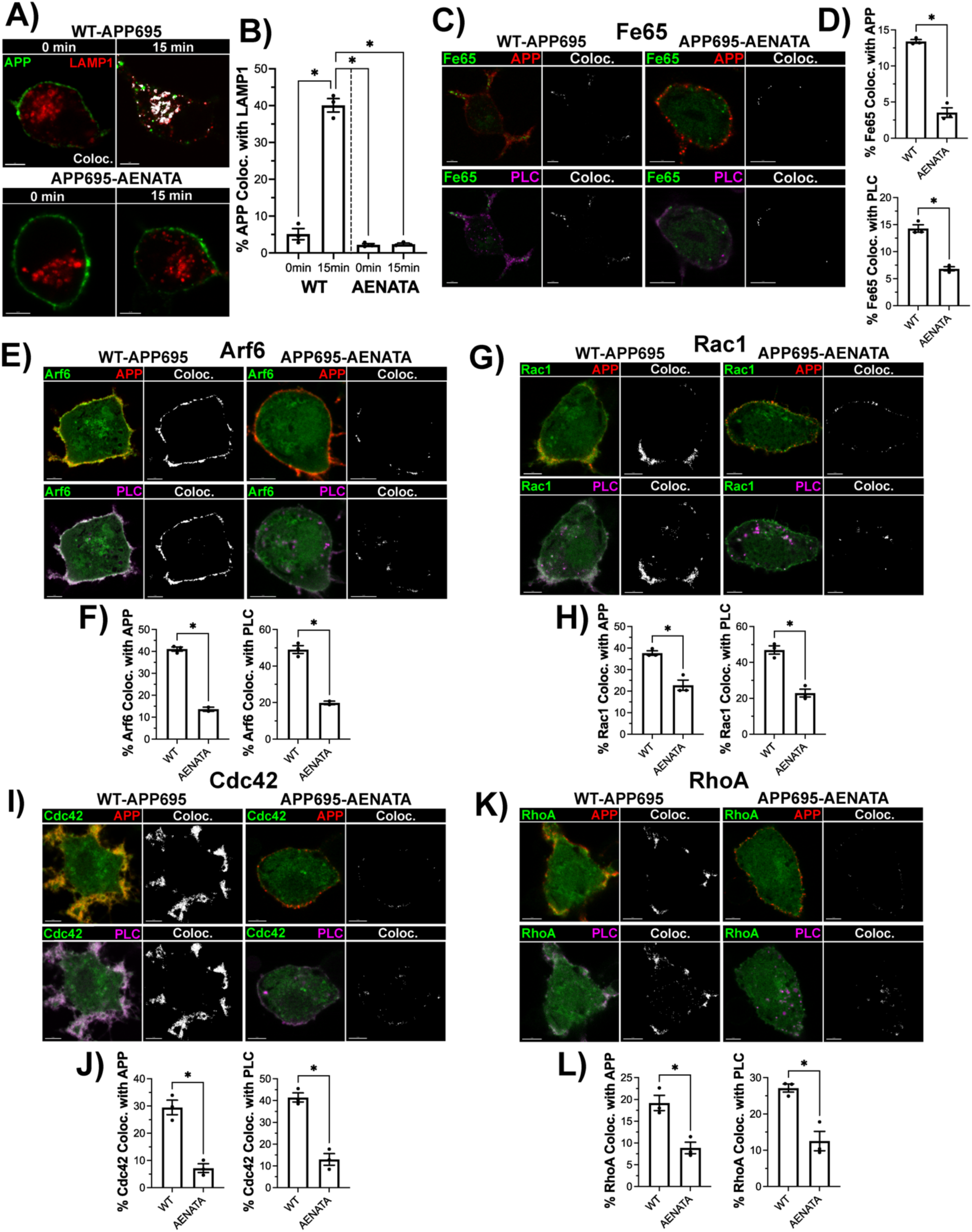
Mutation of the YENPTY site prevents APP macropinocytosis and regulatory protein recruitment. **A)** N2a cells transfected with either WT-APP (top) or APP-AENATA (bottom) and LAMP1-mCh (red). Cells were incubated with tagged anti-APP antibody (green) on ice for 20 minutes, then immediately fixed (0 min) or incubated for 15 minutes. Colocalization was assessed between bound/crosslinked APP and LAMP1 (white pixels). **B)** Quantification of the mean % of APP colocalized with LAMP1 from three replicate experiments (n=3; 15 images per replicate) of cells expressing WT APP or APP-AENATA. Statistical significance was analyzed by a one-way ANOVA with Tukey’s test. **C)** N2a cells transfected with WT-APP or APP-AENATA, Fe65-EGFP (green), and PLC8PH-mRFP (magenta). APP was bound/crosslinked by incubation with a tagged APP antibody (red) while on ice, then incubated for 30 seconds. Colocalization was measured between Fe65 and bound/crosslinked WT-APP or APP-AENATA, as well as between Fe65 and PLC8PH (white pixels). **D)** Quantification of the mean % of Fe65 colocalized with either bound/crosslinked APP (top) or PLC8PH (bottom) in N2a cells expressing WT-APP or APP-AENATA. Data presented comes from three replicate experiments (n=3; 15 images per replicate) of cells expressing WT APP or APP-AENATA. Significant differences between cells expressing WT or AENATA mutant APP was analyzed by a two-tailed unpaired t-test. This experiment was repeated with N2a cells expressing **E/F)** Arf6-GFP, **G/H)** Rac1-GFP, **I/J)** Cdc42-GFP, and **K/L)** RhoA-GFP. *Data is presented as mean ± SEM. * p<0.05; Scale bar = 5 μm*.

### Recruitment of Fe65 and Arf6 to bound/crosslinked APP and membrane ruffles is reduced in response to Arf6 inhibition

To examine if the recruitment of each of these proteins was sequential, we inhibited each GTPase and examined the recruitment of each of the regulatory proteins. N2a cells were transduced with APP695, PLC8PH-mRFP and Fe65-EFGP or Arf6-EGFP. Transduced cells were incubated with inhibitors for Arf6 (NAV-2729), Rac1 (EHT 1864), Cdc42 (ML 141) or RhoA (Rhosin) (see methods). Cells were then incubated with fluorescently labeled APP antibodies on ice for 20 minutes and then incubated at 37°C for 30 seconds, fixed and imaged. Based on studies which have demonstrated that Fe65 acts as a scaffolding protein and recruits Arf6 and Rac1 for actin polymerization [31,32], we predicted that treatment with any GTPase inhibitors tested to have no effect on recruitment. Surprisingly, Arf6 inhibition did significantly reduce the percent of Fe65 colocalized with APP (-8.060 ± 1.392; p=0.001) and PLC8PH (-12.01 ± 1.987; p<0.001) compared to DMSO-treated vehicle control, with no significant reductions being observed with Rac1, Cdc42 or RhoA inhibition (Figure 6A,B).

**Figure 6.**
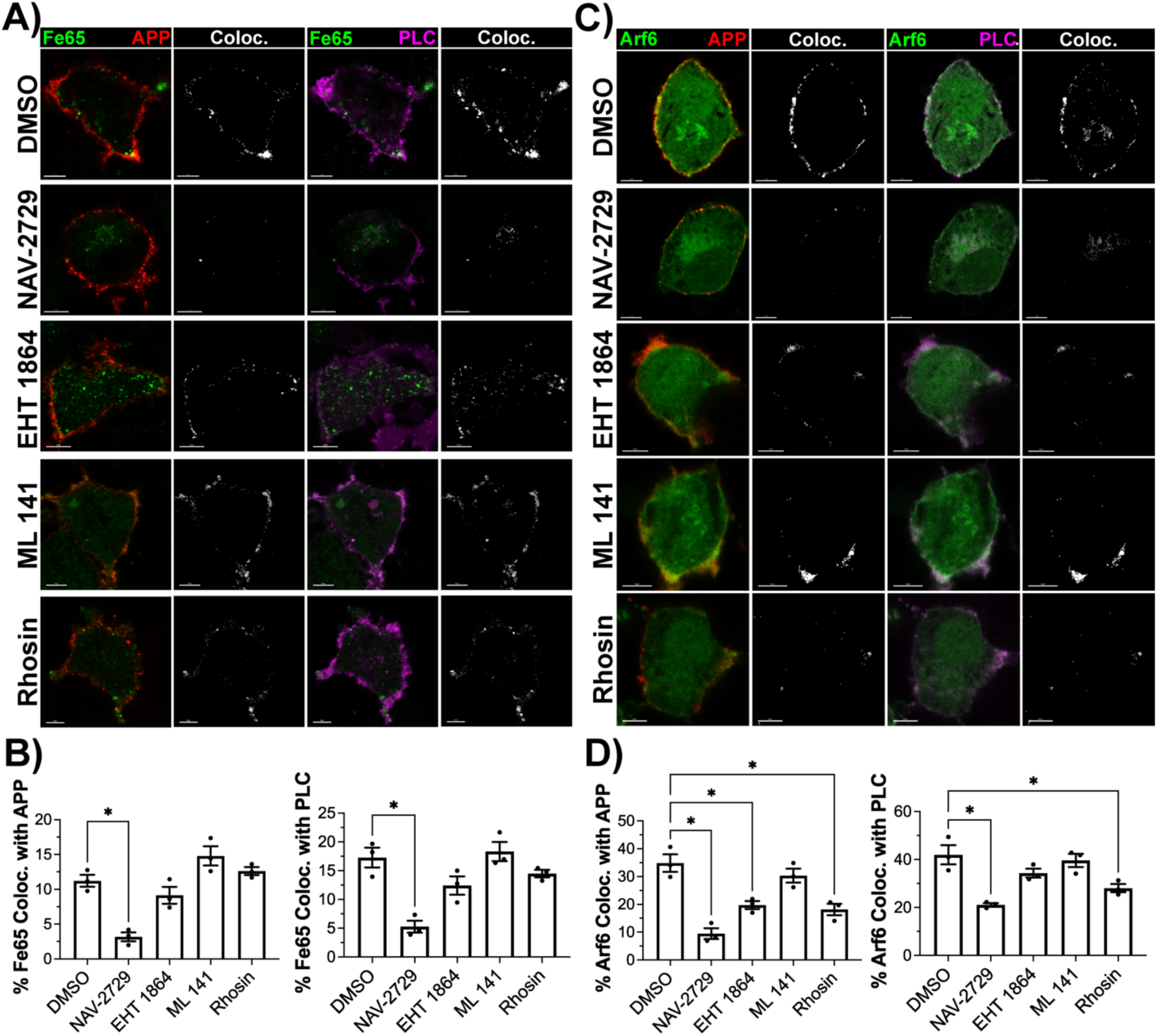
Effects of GTPase inhibition on the recruitment of Fe65 and Arf6 to bound/crosslinked APP and membrane ruffles at 30 seconds. **A)** N2a cells transfected with Fe65-EGFP (green), PLC8PH-mRFP (magenta) and APP695. Cells were treated with 0.1% DMSO, 5μM NAV-2729, 10μM EHT 1864, 10μM ML 141, and 35μM Rhosin. After treatment, cells were incubated with tagged anti-APP antibodies (red) on ice and then fixed following an incubation for 30 seconds. Colocalization was assessed between Fe65 and antibody bound/crosslinked APP or PLC8PH (white pixels). **B)** Quantification of the mean % of Fe65 colocalized with APP (left) or PLC8PH (right) from three replicate experiments (n=3; 15 images per replicate). **C)** N2a cells transfected with Arf6-EGFP (green), PLC8PH-mRFP (magenta) and APP695. Cells were treated with 0.1% DMSO, 5μM NAV-2729, 10μM EHT 1864, 10μM ML 141, and 35μM Rhosin. After treatment, cells were incubated with tagged anti-APP antibodies (red) on ice and then fixed following a 30 second incubation. Colocalization was assessed between Arf6 and bound/crosslinked APP or PLC8PH (white pixels). **D)** Quantification of the mean % of Arf6 colocalized with APP (left) or PLC8PH (right) from three replicate experiments (n=3; 15 images per replicate). *Significant changes in the percentage colocalized between treatments were assessed by a one-way ANOVA with Tukey’s test*. *Data is presented as mean* ± *SEM. *p<0.05; Scale bar = 5μm*.

With Arf6 previously shown to mediate the recruitment and activity of Rac1 in macropinocytosis [25], we hypothesized it acts upstream of the RhoGTPases and predicted only the inhibition of Arf6 itself to result in changes to its recruitment to bound/crosslinked APP and membrane ruffles. As predicted, Arf6 inhibition reduced both its colocalization with APP (-25.35 ± 3.525; p<0.001) and PLC8PH (-20.88 ± 3.506; p<0.001) compared to DMSO vehicle control (Figure 6C,D). However, Rac1 inhibition (-15.10 ± 3.252; p=0.006) and RhoA inhibition (-16.67 ± 3.252; p=0.003) also reduced the percent of Arf6 colocalized with bound/crosslinked APP (Figure 6D), albeit to a lesser extent than Arf6 inhibition. Significant reduction in Arf6 within PLC8PH labelled membrane ruffles was also observed with Arf6 inhibition (-20.88 ± 3.506; p<0.001; Figure 6C,D) and RhoA inhibition (-13.92 ± 3.506; p=0.02; Figure 6C,D).

### The recruitment of the RhoGTPases Rac1, Cdc42 and RhoA to bound/crosslinked APP and membrane ruffles demonstrate differing responses to GTPase inhibition

As above, N2a cells were transduced with APP695, PLC8PH-mCRFP and each of Rac1, Cdc42 or RhoA fused to EGFP and then incubated with the GTPase inhibitors used in the previous experiment (see methods). Cells were then incubated with fluorescently labeled APP antibodies on ice for 20 minutes and then incubated at 37°C for 30 seconds, fixed and imaged. Examining the effects of GTPase inhibition on Rac1, Cdc42 and RhoA recruitment, we found Rac1 colocalization with bound/crosslinked APP was significantly reduced following treatment only with Arf6 inhibitor treatment compared to vehicle control (-30.33 ± 5.503; p<0.001; Figure 7A,B). This reduction was also observed looking at membrane ruffles, with only Arf6 inhibition having a significant effect on the colocalization between Rac1 and PLC8PH (-27.71 ± 5.713; p=0.002; Figure 7A,B). Examining the recruitment of Cdc42, Arf6 inhibition (-27.46 ± 3.679; p<0.001), Rac1 inhibition (-17.68 ± 3.679; p=0.005), and RhoA inhibition (-17.53 ± 3.679; p=0.005) all demonstrated significant reduction in Cdc42-APP colocalization (Figure 7C,D). However, only Arf6 inhibition significantly reduced the colocalization of Cdc42 with PLC8PH (-23.32 ± 4.363; p=0.002; Figure 7C,D). RhoA colocalization with either bound/crosslinked APP or PLC8PH was significantly reduced compared to vehicle control in response to treatment with all of the inhibitors examined (p<0.01; Figure 7E,F).

**Figure 7.**
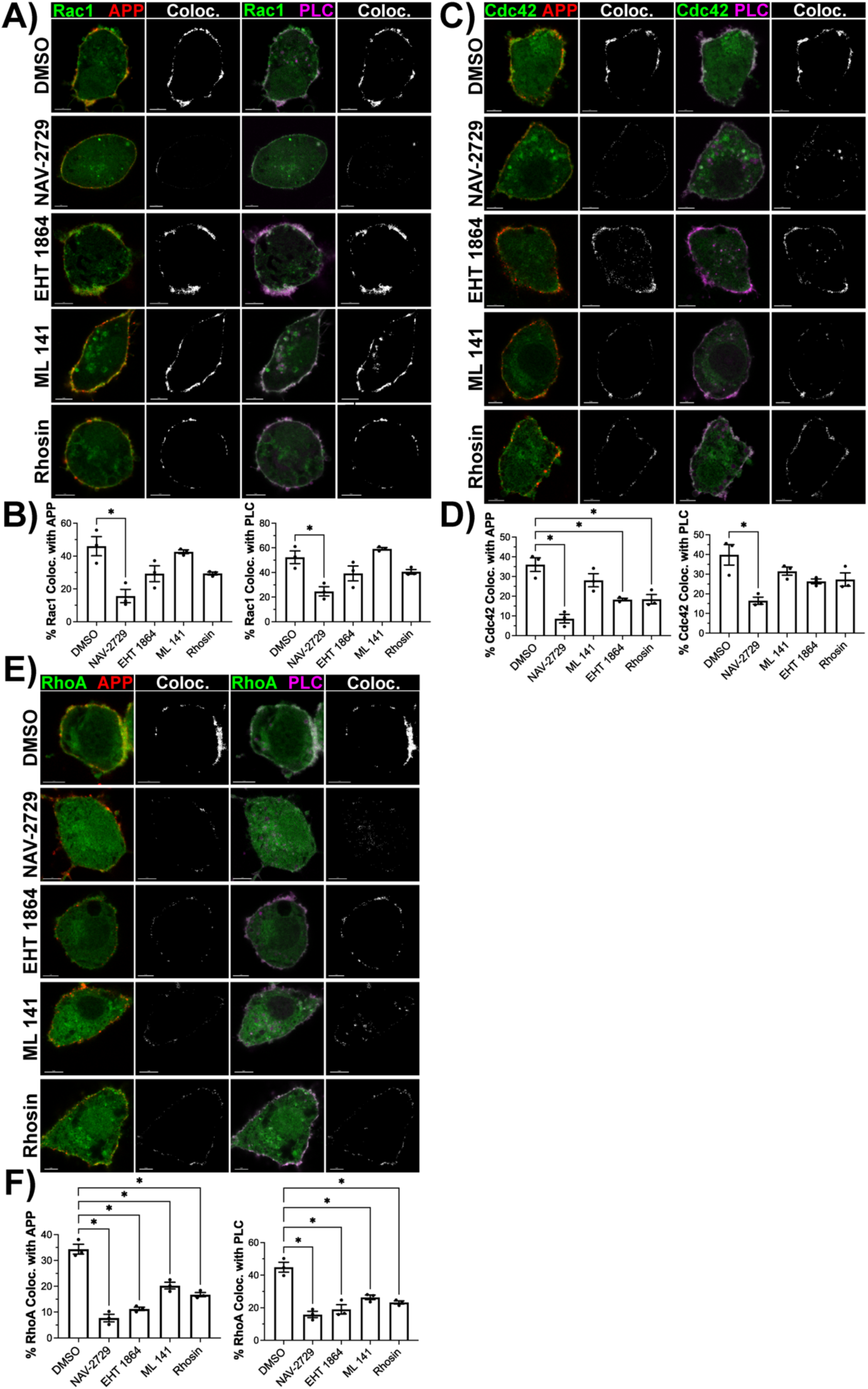
Effects of GTPase inhibition on the recruitment of the RhoGTPases Rac1, Cdc42 and RhoA to bound/crosslinked APP and membrane ruffles at 30 seconds. **A)** N2a cells transfected with Rac1-EGFP (green), PLC8PH-mRFP (magenta) and APP695. Cells were treated with 0.1% DMSO, 5μM NAV-2729, 10μM EHT 1864, 10μM ML 141, and 35μM Rhosin. After treatment, cells were incubated with tagged anti-APP antibodies (red) on ice and then fixed following a 30 second incubation. Colocalization was assessed between Rac1 and bound/crosslinked APP or PLC8PH (white pixels). **B)** Quantification of the mean % of Rac1 colocalized with APP (left) or PLC8PH (right) from three replicate experiments (n=3; 15 images per replicate). **C)** N2a cells transfected with Cdc42-EGFP (green), PLC8PH-mRFP (magenta) and APP695. Cells were treated as described in A), then incubated with tagged anti-APP antibodies (red) on ice and fixed following a 30 second incubation. Colocalization was assessed between Cdc42 and bound/crosslinked APP or PLC8PH (white pixels). **D)** Quantification of the mean % of Cdc42 colocalized with APP (left) or PLC8PH (right) from three replicate experiments (n=3; 15 images per replicate). **E)** N2a cells transfected with RhoA-EGFP (green), PLC8PH-mRFP (magenta) and APP695. Cells were treated as described in A), then incubated with tagged anti-APP antibodies (red) on ice and then fixed following an incubation for 30 seconds. Colocalization was assessed between RhoA and bound/crosslinked APP or PLC8PH (white pixels). **F)** Quantification of the mean % of RhoA colocalized with APP (left) or PLC8PH (right) from three replicate experiments (n=3; 15 images per replicate). *Significant changes in the percentage colocalized between treatments were assessed by a one-way ANOVA with Tukey’s test*. *Data is presented as mean* ± *SEM. *p<0.05; Scale bar = 5μm*.

To assess if any of the inhibitors used produced off target effects on a non-targeted GTPase, we examined the effects of each inhibitor at the concentrations used above on basal Rac1, Cdc42 and RhoA activity using Rac1, Cdc42 and RhoA G-LISA assays (Cytoskeleton) in untransfected N2a cells. Only treatment with EHT 1864 resulted in significantly reduced GTP-bound Rac1 (p<0.05; Figure 8A). Likewise, only treatment with the inhibitor ML 141 resulted in a significant reduction in the amount of GTP-bound Cdc42 (p<0.05; Figure 8B). Levels of RhoA-GTP were observed to only be significantly reduced in response to treatment with the inhibitor Rhosin (p<0.05; Figure 8C).

**Figure 8.**
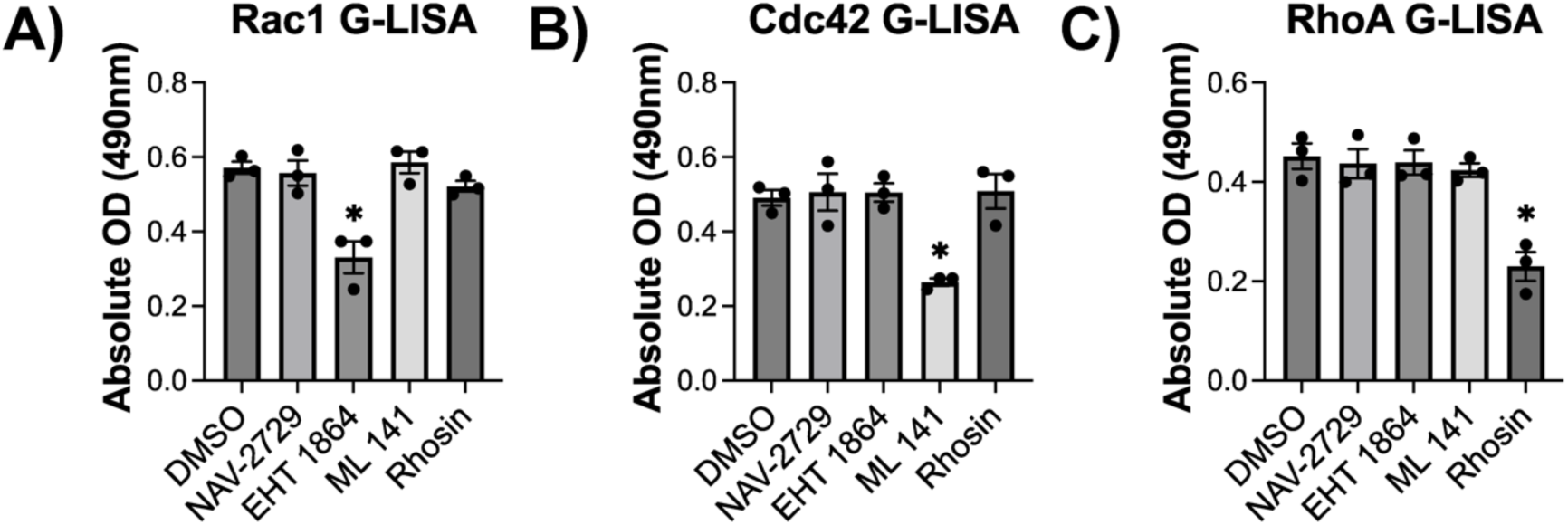
Small molecule GTPase inhibitors selectively decrease basal active GTP-bound protein levels of its target GTPase. **A)** G-LISA assay for Rac1-GTP from N2a cells treated with 0.1% DMSO, 5μM NAV-2729, 10μM EHT 1864, 10μM ML 141, and 35μM Rhosin as used in the experiments above. After treatment, lysates were immediately collected and used for Rac1 G-LISA activation assays. To compare the relative amount of active GTP-bound Rac1 in each treatment condition, the absolute optical density (OD) was measured from each condition’s lysate loaded into G-LISA assay wells. Three different lysate samples were collected for each condition from different biological samples (n=3), and the G-LISA assay was performed in technical triplicates. Significant differences from untreated samples were calculated by a one-way ANOVA with Tukey’s test. This same experiment and analysis was performed for **B)** Cdc42-GTP levels, and **C)** RhoA-GTP levels. *Data is presented as mean* ± *SEM. *p<0.05*.

## Discussion

The internalization of APP is a key step in the production of Aβ. Utilizing an antibody directed towards the N-terminal domain of APP to bind/crosslink APP and trigger macropinocytosis, we have demonstrated the direct recruitment of Fe65, Arf6, Rac1, Cdc42 and RhoA to bound/crosslinked APP and PLC8PH containing membrane ruffles. Additionally, using GTPase inhibitors and mutation of the YEPNTY sequence of APP, we demonstrated the sequential nature of this regulatory cascade. Our results suggest that Fe65 is recruited to APP, which then mediates the recruitment of Arf6, followed by downstream recruitment and activation of Rac1, Cdc42 and RhoA. To our knowledge, this is the first evidence which demonstrates the presence of a regulatory signalling cascade mediating the macropinocytosis of APP, reinforcing APP’s role as a receptor for the stimulation of macropinocytosis following binding to its ectodomain or its dimerization. These results support our prior observations which utilized dominant negative mutant Arf6, and both small molecule inhibition and siRNA-mediated knockdown of Rac1, RhoA and Cdc42 [14,34], which demonstrated that the activity of these proteins was required for the rapid internalization of APP to lysosomes.

Previous studies have demonstrated that inhibition of Rac1 and ROCKII, a downstream effector of RhoA, decreased Aβ40 and Aβ42 production [28,29]. In both studies, the mechanism underlying the reduction in Aβ as a result of Rac1 and ROCKII inhibition was suggested to be due to altered trafficking of APP and BACE or γ-secretase leading to decreased interactions [28,29]. While they both provided suggestions of the underlying mechanism, neither study directly investigated the exact trafficking mechanisms involved, how they might have been altered because of GTPase inhibition, and how this impacted trafficking to an end-target. The findings of this study and our previous work would suggest that reduced APP trafficking to lysosomes through macropinocytosis is the mechanism underlying these previous observations. However, how APP macropinocytosis and the amyloidogenic processing of APP are connected in normal physiologically and AD pathophysiology is still unclear. Alterations in the co-trafficking of BACE and γ-secretase could provide one explanation linking the two, as suggested by the study discussed above investigating ROCKII inhibition [29]. Understanding why APP binding or crosslinking results in its macropinocytosis, how it contributes to the amyloidogenic processing of APP, and if this mechanism produces pathogenic production of Aβ in AD would provide new insights into the role of APP in AD pathogenesis.

Both APP and macropinocytosis have been connected to growth cone function and axonal extension and turning and are broadly connected to synaptic plasticity. APP has been shown to function in the formation of synapses. Knock-outs of APP have demonstrated that it plays an important role long-term potentiation (LTP) and synaptic plasticity in the adult mouse brain [38]. On cellular level, it has been observed that APP is enriched in both growth cones and synapses [39,40]. Here, it has been shown to regulate neurite and growth cone formation through interactions with various stimulating factors such as neural growth factor (NGF) [39] and Netrin-1 [41]. Our observations supporting the regulatory function of Fe65 and Arf6 in APP macropinocytosis may provide insights linking these previous studies. Arf6 has been shown to mediate neurite outgrowth through Fe65 [31] and Fe65 itself is enriched in growth cones and synapses [40]. Furthermore, regulatory roles of RhoGTPases Rac1, Cdc42 and RhoA in neurite outgrowth and growth cone formation have also been demonstrated [42]. However, direct investigation of how APP mediates its role in growth cone activity and synapse formation has not yet been investigated.

Although APP was sequenced in 1987 [22], its function remains unknown. APP was suggested to act as a cell surface receptor since its identification, and studies of its structure have revealed domains that can mediate ligand binding and formation of APP dimers [35]. The results demonstrated in this study, along with our current and previous findings, could show that APP acts as a cell surface receptor stimulating its macropinocytosis. This is similar to growth factor receptors, where ligand binding frequently results in their internalization from the membrane by macropinocytosis [36]. In fact, the homodimerization of growth factor receptors in response to ligand binding and subsequent C-terminal phosphorylation has been observed in several growth-factor-induced macropinocytosis mechanisms [18]. This model of APP-stimulated macropinocytosis could further explain the previously discussed links between APP and macropinocytosis in growth cones.

Given the lack of a putative APP ligand, antibody-driven crosslinking of APP remains our best experimental model for studying this. Supporting our findings using this approach, antibody-mediated crosslinking of APP has been utilized by several other labs which also demonstrated increased endocytosis of APP as a result [23]. The presence of the GFLD in the E1 domain of APP [35] could also be argued to support this proposed APP-stimulated macropinocytosis model. This domain demonstrates a high degree of structural similarity with growth-factor receptors, and the broader E1 domain has been demonstrated to bind to a number of extracellular proteins [35,43]. This includes Reelin and NGF [44], which has been shown to stimulate macropinocytosis within growth cones [45]. Further, YENPTY and similar sequences have been shown to regulate the recruitment of macropinocytosis regulatory machinery to receptors which stimulate macropinocytosis [18,46].

However, it should be noted that dimerization of APP, and phosphorylation of tyrosine residues within the YENPTY sequence have been implicated in other endocytic mechanisms including clathrin-mediated endocytosis (CME). Phosphorylation of the first tyrosine of the YENPTY sequence was required for the immunoprecipitation of APP with adaptor protein-2 (AP2) and clathrin [47]. While our original study investigating the trafficking of crosslinked APP revealed rapid and direct trafficking to the lysosome, crosslinked APP was also observed in Rab5-labelled early endosomes [13]. It is worth noting that internalization by both CME and macropinocytosis has been observed in many growth-factor receptors [48], and could occur with APP. Many questions still remain regarding the function and physiological role of APP and how changes to its function contribute to AD pathogenesis. Gaining a complete understanding of its endocytic trafficking mechanisms, such as macropinocytosis, could lead to new insights on its functions and what leads to the overproduction of Aβ in AD.

## Conclusion

Here, we have demonstrated the direct recruitment of the macropinocytosis regulatory proteins Arf6, Rac1, Cdc42, and RhoA to crosslinked APP. Further we have provided evidence for Fe65 acting as a scaffolding protein for the recruitment of these proteins to crosslinked APP. Together, these observations demonstrate a regulatory cascade works to govern the internalization of APP to lysosomes by macropinocytosis. Since GTPases are readily targeted by pharmacological inhibitors, this cascade could be explored as a potential novel avenue for therapies aimed at reducing Aβ production by reducing the trafficking of APP from the cell surface to lysosomes.

## Materials and Methods

### Antibodies and Reagents

The antibodies purchased were: mouse N-terminal anti-APP antibody (mAbP2-1, OMA1-03132, 1:100, Invitrogen, MA, USA), mouse Aβ region anti-APP (6E10, 803001, 1:100, Biolegend, CA, USA) and rabbit anti-NCAM1 antibody (EPR2187, ab220360, 1:00, Abcam, MA, USA). Zenon Alexa Fluor 647 Mouse IgG1 Labeling Kit (Z25008) and Zenon Alexa Fluor 647 Rabbit IgG Labelling Kit (Z25308) were purchased from Invitrogen (MA, USA). The Arf6 inhibitor NAV 2729 (SML2238), Rac1 inhibitor EHT 1864 (E1657), Cdc42 inhibitor (SML0407), and Wortmannin (681675) were purchased from Sigma-Aldrich (MO, USA). The RhoA selective inhibitor Rhosin hydrochloride (5003) and ethylisopropyl amiloride (EIPA; 3378) were purchased from Tocris (5003; Bristol, UK). Pitstop2 was purchased from abcam (ab120687; MA, USA). Alexa488 conjugated transferrin was purchased from Invitrogen (T13342, MA, USA). Neuro-2a (N2a) mouse neuroblastoma cells were acquired from ATTCC (CCL-131, VA, USA). The reagents: Dulbecco’s phosphate-buffered saline (DPBS, 14190144), Hank’s balanced salt solution (HBSS, 14025092) and trypsin-EDTA (25200056) were purchased from Gibco (CA, USA); fetal bovine serum (FBS, 090-150) from Wisent (QC, Canada); paraformaldehyde (PFA, 043368.9M) from Thermo Scientific (MA, USA); dimethyl sulfoxide (DMSO, D8418) from Sigma-Aldrich (MO, USA). G-LISA kits for GTP-bound Rac1 (BK128), Cdc42 (BK127) and RhoA (BK124) were all purchased from Cytoskeleton (CO, USA).

### DNA Constructs

The untagged APP695 construct was kindly provided by Dr. Jane Rylett (Robarts Research Institute, ON, Canada). APP695-AENATA (pCAX-APP-AENATA) was a gift from Dennis Selkoe and Tracy Young-Pearse (Addgene plasmid #30144). LAMP1 fused to mCherry (LAMP1-mCh) construct was described in a previous study by our lab [49]. Arf6-EGFP was a generous gift from Dr. Susan Meakin (Western University, ON, Canada). Rac1-EGFP (pcDNA3.1-EGFP-Rac1(wt)) was a gift from Klaus Hahn (Addgene plasmid #13719). Cdc42-EGFP (pcDNA3.1-EGFP-Cdc42-wt) and RhoA-EGFP (pcDNA3.1-EGFP-RhoA-wt) was a gift from Gary Bokoch (Addgene plasmid #12975 and #12965). PLCδPH-mRFP was a generous gift from Dr. Bryan Heit (Western University, ON, Canada). Fe65-EGFP was a construct produced by VectorBuilder (IL, USA).

### Cell Culture and Transfection

Neuro2a (N2a) cells were cultured in MEM containing 10% FBS in a 25 cm^2^ flask (130189, Thermo Scientific, MA, USA) at 5% CO2 at 37°C. The cells were passaged by trypsinization every 3-4 days. Once plated for experiments, cells were transiently transfected using Lipofectamine 2000 (11668019, Invitrogen, MA, USA) according to the manufacturer’s protocol. After 24 hours post-transfection, N2a cells were differentiated by serum withdrawal (i.e., serum-free MEM) for 18h. This differentiation method was chosen as serum-withdrawal has been demonstrated to induce neurite outgrowth and increased expression of neuronal protein markers such as NeuN [50,51]. Following differentiation, cells were used for experiments as described below.

### Inhibitor Treatments

Following differentiation, cells were treated with inhibitors in fresh serum-free media. Small molecule GTPase inhibitors, EIPA and Pitstop2 were reconstituted from the manufacturer in DMSO. As a vehicle control, 0.1% DMSO (v/v) in serum-free medium was used. For experiments examining the recruitment of the regulatory proteins of interest to crosslinked APP the following concentrations and incubation times were used: 5 μM NAV-2729 (3 hrs), 10 μM EHT 1864 (18 hrs), 10 μM ML 141, 35 μM Rhosin (3 hrs), and 0.1% DMSO (18 hrs). Treatment with EHT and DMSO occurred overnight during differentiation of N2a cells, and the remaining treatments were performed following differentiation. EHT 1864 and ML 141 concentrations were used in our previous study [34] and confirmed to inhibit rapid APP internalization to lysosomes in this study (Figure S2). NAV-2729 and Rhosin treatments were chosen based on previous literature using these inhibitors [52,53], and a dose titration was performed for each to determine the doses used in this current study to inhibit rapid internalization of APP to lysosomes (Figure S2).

Amiloride and amiloride-based compounds like EIPA are commonly used as inhibitors of macropinocytosis by reducing the activity of Rac1 and Cdc42 through local plasma membrane pH changes produced by blocking the activity of Na+/H+ exchangers [54]. To assess its effects on APP internalization to lysosomes, cells were treated with 10 μM EIPA or 0.1% DMSO for 1 hour in serum-free media. Pitstop2 was used to inhibit CME, as it has been shown to inhibit clathrin-coated pit formation [55]. This was chosen because dynamin inhibition using Dynasore has off-target effects and been shown to result in the inhibition of macropinocytosis [56]. Pitstop2 treatment was performed directly before crosslinking APP by incubating N2a cells with 20 μM Pitstop2 for 5 minutes. This treatment dose and timing was recommended by the manufacturer and validated by demonstrating reduced Alexa488 conjugated transferrin uptake in N2a cells following Pitstop2 treatment as described (Figure S3).

### Antibody-mediated cell surface APP binding/crosslinking

As performed in our previous studies on APP macropinocytosis [13,14,34], the binding and/or crosslinking of APP was used to stimulate its internalization by macropinocytosis. For cell surface APP binding/crosslinking experiments, N2a cells were seeded at a density of 1.20 x 10^5^ cells in a 35mm dish with a 14mm glass-bottom (P35G-1.5-14-C, MatTek, MA, USA). Anti-APP (Invitrogen, MA, USA) was labelled with Alexa Fluor 647 using a Zenon Alexa 647 Mouse IgG1 Labeling Kit (Invitrogen, MA, USA) according to the manufacturer’s instructions. As a negative control, anti-NCAM antibody (Abcam, MA, USA) was labelled with Alexa Fluor 647 using a Zenon Alexa 647 rabbit IgG Labeling Kit (Invitrogen, MA, USA) according to the manufacturer’s instructions and used to label cell surface NCAM. Labelled antibodies were incubated with N2a cells in HBSS on ice for 20 minutes. Previously, our lab has demonstrated incubation with anti-APP antibodies on ice allowed for thorough labelling of cell surface APP but halted the cell’s ability to endocytose, and subsequent removal from ice stimulated the macropinocytosis of antibody bound/crosslinked APP [14]. For a baseline measurement of regulatory protein recruitment, following incubation with antibodies on ice, cells were washed with ice-cold HBSS to remove unbound antibody and fixed on ice for 45 minutes with ice-cold 4% PFA (denoted as 0 min). For the 30-second time point, the cells were taken off the ice, washed once with HBSS pre-warmed to 37 °C to remove unbound antibody and immediately fixed (denoted as 30 sec). For all other time points, after washing, warm HBSS was added to the cells, and they were incubated at 37 °C and 5% CO_2_ at indicated time points before fixing. Fixation for all time points (except 0 min) was performed by adding 4% PFA to cells for 15 minutes at room temperature. Cells selected for imaging had normal morphology and good expression of transfected constructs. All experiments were replicated three times, with at least 10 or 15 cells sampled for each condition at each time point.

### Confocal Microscopy

Imaging was performed using a Leica SP8 confocal microscope and an HC PL APO CS2 63X 1.4 numerical aperture oil immersion lens. The optical section thickness was typically 1 micron. GFP fluorescence was imaged using a 488 nm excitation laser with a 500-550 nm filter set on a HyD hybrid detector. mRFP and mCherry fluorescence were visualized using a 542 nm excitation laser with a 570-620 nm filter on a PMT detector. Alexa Fluor 647 fluorescence was imaged using a 638 nm excitation laser with a 650-700 nm filter set on a HyD hybrid detector. Z-stacks were taken to image the entirety of the cells.

### Data Quantification and Analysis

Colocalization analysis was performed on the z-stack images using Imaris 10.1.0 imaging software (Bitplane, CT, USA). To assess colocalization, I utilized an approach to setting the colocalization threshold of the protein of interest to select only the brightest 0.5% of pixels in the channel of interest, adapted from Lorenzen et al. (2010) to consider signal in an unbiased manner based on the intrinsic property of the image [13]. The threshold adopted for the other channel of interest (in most experiments this is LAMP1, crosslinked APP, or PLC8PH), was selected based on the average pixel intensity that demarcated a clear structure (vesicle or membrane ruffle) to ensure only clear signal was considered in the analysis. For example, to determine the recruitment of Arf6, the colocalization threshold for Arf6 was set to the brightest 0.5% pixels. This allowed for quantification of changes in accumulation of localized Arf6. To assess its colocalization with crosslinked APP, a threshold was set based on the average membrane signal intensity demarked by crosslinked APP. Imaris then generated the percentage colocalized by determining the number of pixels (above the threshold) of the two specified channels. Using this unbiased method of measuring percentage colocalized, we compared the percentage colocalized between timepoints and conditions, all analyzed using the method described above. Graphing and statistical analysis were performed using Prism GraphPad 6.0 using either unpaired two-tailed t-tests or one-way ANOVAs with Tukey’s test. p-values of less than 0.05 were considered to be statistically significant.

### Assessing RhoGTPase activity following APP crosslinking using GLISA

For culturing cells for G-LISA assays, N2a cells were cultured in MEM containing 10% FBS in a 25 cm^2^ flask (130189, Thermo Scientific, MA, USA) at 5% CO2 at 37 °C. The cells were passaged by trypsinization every 3-4 days. One day before transfection, cells were seeded at a density of 5.0 x 10^4^ cells in each well of a 4-well plate. Cells were subsequently transfected with the APP695 construct using lipofectamine 2000. The following day, the cells were serum starved overnight for 18 hours to ensure low basal levels of GTPase activity and induce differentiation. Following serum starvation, cells were placed on ice and a 4-well plate was each treated with anti-NCAM or anti-APP antibodies or with HBSS (no-treatment control). As a positive control, a 4- well plate was treated with EGF. Each 4-well plate was incubated with its respective treatment for 20 minutes on ice then moved to an incubator at 5% CO2, 37 °C for 5 minutes. Following this incubation, plates were immediately moved back to ice, washed two times with ice-cold PBS, and then extracted protein according to manufacturer protocol. G-LISAs were then run according to manufacturer protocol in technical and biological triplicates.

