## Supplemental Figures S1-3 for "Macropinocytosis of amyloid precursor protein requires the adaptor protein Fe65 and the recruitment and activity of Arf6 and the RhoGTPases Rac1, Cdc42 and RhoA"

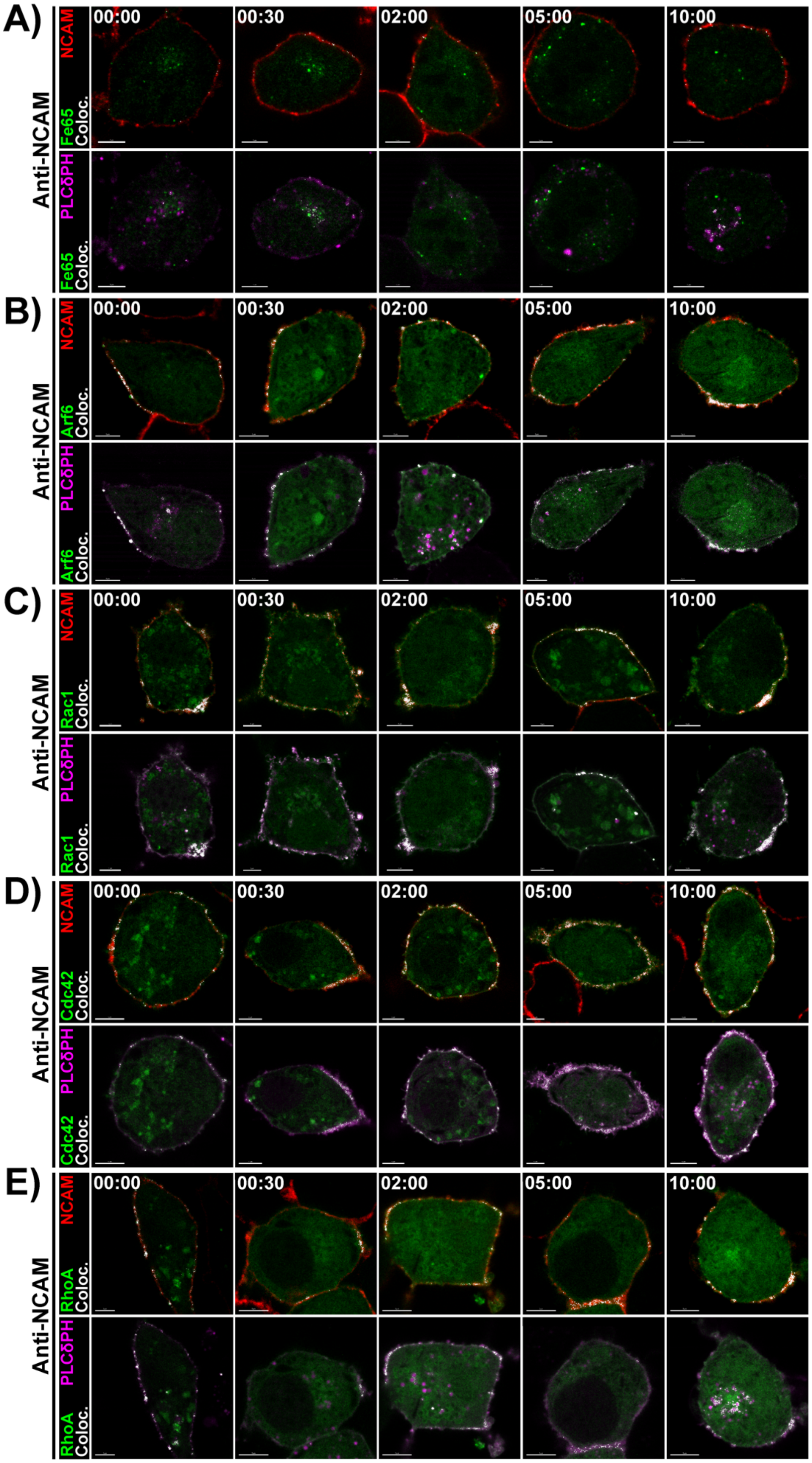
A)
B)
C)
D)
E)
Anti-NCAM
NCAM
Fe65
Coloc.
PLCδPH
Arf6
Rac1
Cdc42
RhoA
00:00
00:30
02:00
05:00
10:00

**Figure S1. Antibody-mediated binding/crosslinking of NCAM negative control for regulatory protein recruitment.**

**A)** N2a cells transfected with Fe65-EGFP (green), PLCδPH-mRFP (magenta) and APP695. Cells were incubated with fluorescent-tagged anti-NCAM antibodies (red) on ice, then immediately fixed on ice as a baseline (00:00) or incubated for 30 seconds (00:30), 2 minutes (02:00), 5 minutes (5:00), and 10 minutes (10:00) and then fixed. Colocalization between Fe65-EGFP and anti-NCAM or PLCδPH-mRFP signal is indicated by white pixels. This experiment was repeated with cells expressing **B)** Arf6-GFP, **C)** Rac1-GFP, **D)** Cdc42-GFP and **E)** RhoA-GFP. *Scale bar = 5µm.*

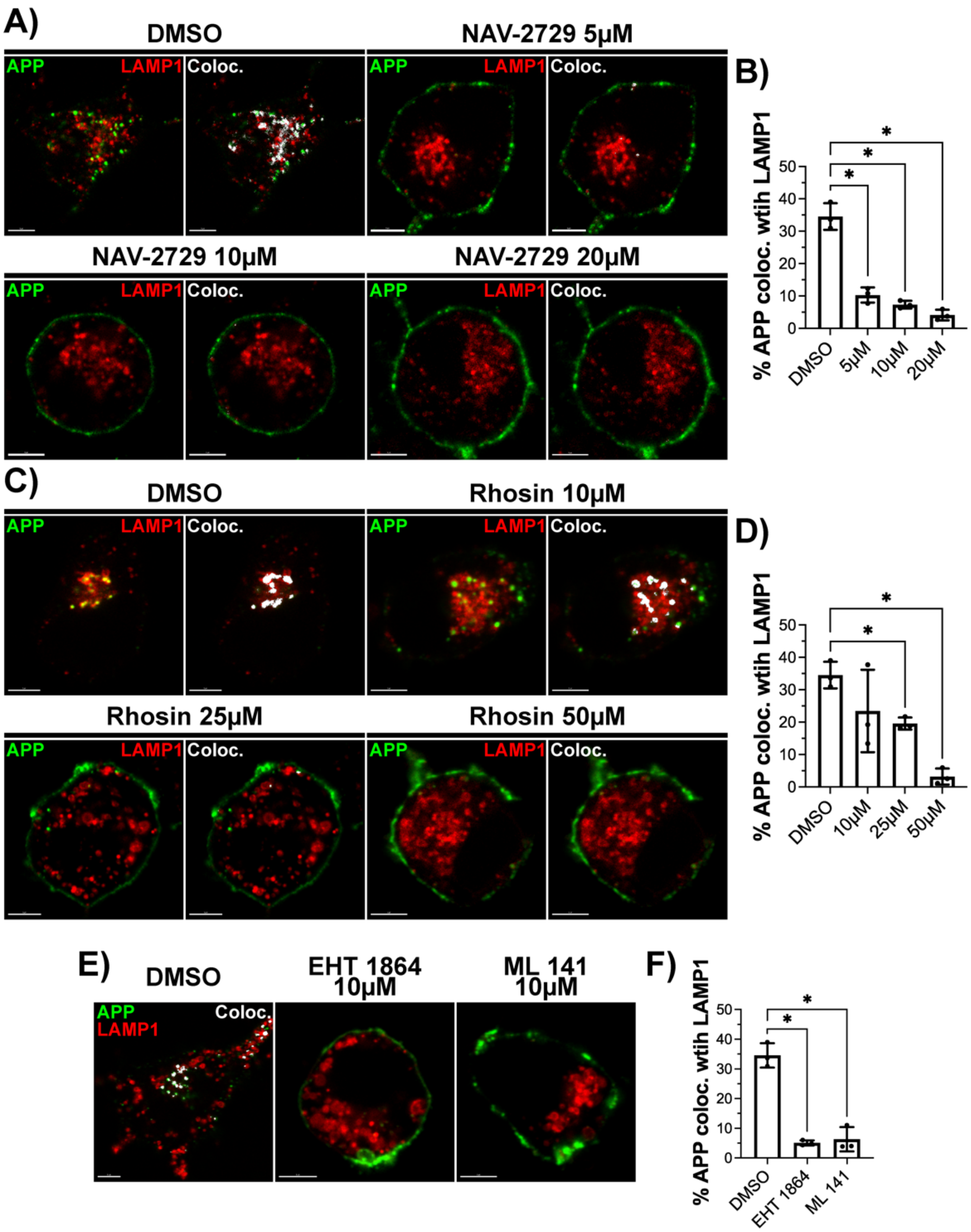


**Figure S2. Inhibition of GTPases using NAV-2729, Rhosin, EHT 1864 and ML 141 reduces rapid internalization of APP to lysosomes.**

**A)** N2a cells transfected with APP695 and LAMP1-mCh (red) that were treated with DMSO, 5µM, 10µM, or 20µM NAV-2729 (Arf6 inhibitor). APP was then bound/crosslinked by fluorescent tagged APP antibodies and imaged after 15 minute incubation. Colocalization was assessed between crosslinked APP and LAMP1 (white pixels). **B)** Quantification of the mean % of APP colocalized with LAMP1 (n=3; 10 images per replicate), with significance calculated by a one-way ANOVA with Tukey's test. **C)** N2a cells transfected with APP695 and LAMP1-mCh (red) that were treated with DMSO, 10μM, 25μM, or 50μM Rhosin (RhoA inhibitor). APP was then bound/crosslinked by fluorescent tagged APP antibodies and imaged after 15 minute incubation. Colocalization was assessed between crosslinked APP and LAMP1 (white pixels). **D)** Quantification of the mean % of APP colocalized with LAMP1 (n=3; 10 images per replicate), with significance calculated by a one-way ANOVA with Tukey's test. **E)** N2a cells transfected with APP695 and LAMP1-mCh (red) that were treated with DMSO, 10μM EHT 1864 (Rac1 inhibitor), or 10μM ML 141 (Cdc42 inhibitor). APP was then bound/crosslinked by fluorescent tagged APP antibodies and imaged after 15 minute incubation. Colocalization was assessed between crosslinked APP and LAMP1 (white pixels). **F)** Quantification of the mean % of APP colocalized with LAMP1 (n=3; 10 images per replicate), with significance calculated by a one-way ANOVA with Tukey's test. *Data is presented as mean ± SEM. * $p<0.05$; Scale bar = 5μm.*

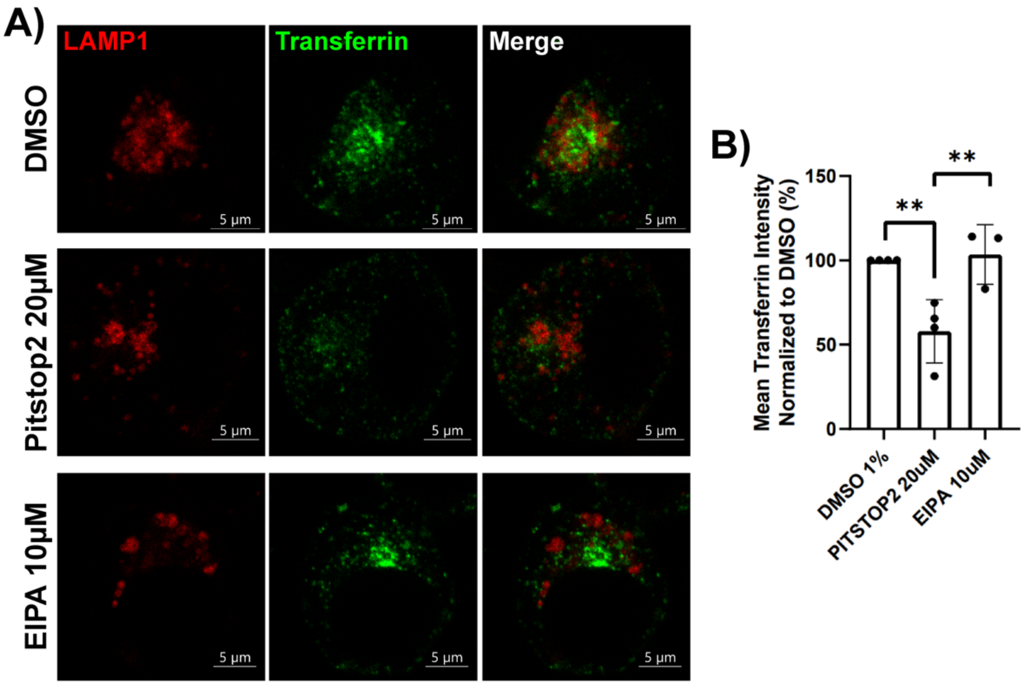


**Figure S3. Pitstop2 inhibition of CME reduces the amount of transferrin internalized in N2a cells.**

**A)** N2a cells transfected with LAMP1-mCh (red) that were treated with DMSO (vehicle control), 20µM Pitstop2 (CME inhibitor), or 10µM EIPA (macropinocytosis inhibitor). Transfected cells were incubated with Alexa488-tagged transferrin on ice for 20 minutes then fixed and imaged after a 15 minute incubation. Imaging settings were kept consistent across images taken and the mean pixel intensity of transferring was measured in each image across all conditions. **B)** Quantification of changes in mean pixel intensity of transferrin signal within N2a cells across treatment conditions (n=3; 10 images per replicate), with significance calculated by a one-way ANOVA with Tukey's test. Mean pixel intensity of transferrin signal was normalized to DMSO vehicle control. Significantly reduction in mean pixel intensity of transferrin signal was observed in response to Pitstop2 treatment indicating inhibition of CME. *Data is presented as mean ± SEM. * $p<0.05$, ** $p<0.01$; Scale bar = 5µm.*
